# Structural basis of lipopolysaccharide translocon assembly mediated by the small lipoprotein LptM

**DOI:** 10.1101/2025.01.03.631273

**Authors:** Ryoji Miyazaki, Mai Kimoto, Hidetaka Kohga, Tomoya Tsukazaki

**Author notes:** corresponding: Correspondence (T. T.) (R. M.). equally contributed (Ryoji Miyazaki, Mai Kimoto, Hidetaka Kohga).

## Abstract

Gram-negative bacteria possess an outer membrane (OM) that acts as a barrier against toxic compounds. Lipopolysaccharide (LPS) in the outer leaflet of the OM is crucial for barrier function. After its synthesis in the cytoplasm and translocation to the periplasm, LPS is transported to the OM by the LPS transport system. The LPS translocon, composed of an OM protein LptD and a lipoprotein LptE, mediates LPS assembly into the OM. Recently, the small lipoprotein LptM (YifL) was identified as a novel LptD/E-interactor that facilitates LptD maturation. However, its mechanism remains unclear. Here, we investigated the detailed interaction between LptM and LptD. We found that LptM interacts with the folded LptD intermediate at the late stage of its maturation. Mutational analyses demonstrated that the N-terminal conserved region (C_20_GLKGPLYF_28_) of LptM is essential for its function. Cryo-EM structural analysis of the *E. coli* LptD/E/M complex, combined with biochemical analyses, revealed the molecular basis of the LptM–LptD interaction and its functional importance. Thus, we propose that LptM functions as a “barrel-rivet,” stabilizing LptD for its proper assembly.

## Introduction

Gram-negative bacteria possess two membranes: the inner (cytoplasmic) membrane (IM) and the outer membrane (OM) (Figure S1). The periplasm is a space between these membranes and contains the peptidoglycan cell wall. The OM faces an extracellular environment and acts as a selective permeability barrier, excluding large hydrophobic compounds such as macrolide antibiotics (Nikaido, 2003). The functionality and structural integrity of the OM rely on various β-barrel outer membrane proteins (OMPs) and lipopolysaccharides (LPSs) at its outer leaflet (Sperandeo *et al*, 2017; Benn *et al*, 2024). The assembly of OMPs and the transport of LPS into the OM are mediated by the β-barrel assembly machinery (BAM) and the LPS transport (Lpt) complexes, respectively. These complexes are essential for cell viability, making them attractive targets for novel antibiotics.

In *Escherichia coli* (*E. coli*), the BAM complex consists of BamA, an essential OMP, and four OM lipoproteins, BamB, BamC, BamD, and BamE (Bakelar *et al*, 2016; Gu *et al*, 2016). Following synthesis in the cytoplasm and membrane-translocation via the SecY/E/G translocon, SecA, SecD/F, and PpiD/YfgM (Tsukazaki, 2018; Miyazaki *et al*, 2022), nascent OMPs are delivered to the BAM complex in the OM by periplasmic chaperones such as SurA, DegP, and Skp, where they are assembled and integrated into the OM (Devlin & Fleming, 2024; Combs & Silhavy, 2024). The mechanism of BAM-mediated OMP assembly has been well studied. Cryogenic electron microscopy (cryo-EM) analyses have resolved many structures of the BAM complex in various states, including substrate proteins, inhibitors, and associated factors (Tomasek *et al*, 2020; Wu *et al*, 2021; Doyle *et al*, 2022; Shen *et al*, 2023; Kaur *et al*, 2021; Sun *et al*, 2024; Fenn *et al*, 2024). These structural studies have provided insights into the OMP assembly process. After being delivered to the BAM complex, the C-terminal regions of substrate OMPs containing a “β- signal” are recognized by the C-terminal strand of the BamA β-barrel domain and BamD (Doyle *et al*, 2022; Germany *et al*, 2023). The β-barrel domain of the OMP then forms a β-barrel and is integrated into the OM through the lateral gate of BamA. After release from the BAM complex, the substrate OMP completes β- barrel closure, achieving its maturation. Recently, biochemical analyses have elucidated the functional roles of the accessory Bam factors BamB, BamC, and BamE, further refining our understanding of the BAM complex (Gunasinghe *et al*, 2018; Thewasano *et al*, 2023).

LPS is synthesized at the inner leaflet of the IM, flipped to the periplasmic side of the IM by the ATP-binding cassette transporter MsbA, and transported from the IM to the OM by the Lpt complex (Okuda *et al*, 2016). The LptB_2_/F/G complex at the IM utilizes ATP hydrolysis energy to extract LPS from the IM (Li *et al*, 2019). Once extracted, LPS is delivered to the OM along a continuous bridge formed by LptF, LptC, the periplasmic protein LptA, and the OMP LptD (Sherman *et al*, 2018; Törk *et al*, 2023). LptD, in complex with the OM lipoprotein LptE, forms the LPS translocon, which transports LPS from the bridge to the outer leaflet of the OM (Qiao *et al*, 2014; Dong *et al*, 2014; Botos *et al*, 2016). LptD is an essential OMP composed of a C-terminal transmembrane β-barrel domain and an N-terminal soluble β-taco domain. In its mature form (LptD^NC^), two intramolecular disulfide bonds form between nonconsecutive Cys pairs (C31-C724 and C173-C725) in LptD, connecting the C-terminal region of the β-barrel domain to the taco domain (Figure 1A) (Qiao *et al*, 2014; Botos *et al*, 2016). In contrast, an LptD intermediate (LptD^C^) is first generated, in which two disulfide bonds are formed between consecutive Cys pairs (C31-C173 and C724- C725) (Figure 1A). During its maturation, the isomerization of disulfide bonds from LptD^C^ to LptD^NC^ is facilitated by LptE when the LptD intermediate is assembled on the BAM complex (Chng *et al*, 2012; Narita *et al*, 2013; Lee *et al*, 2016). The maturation process of LptD has been well studied, resulting in the identification of several proteins involved in this process, such as BepA.

**Figure 1.**
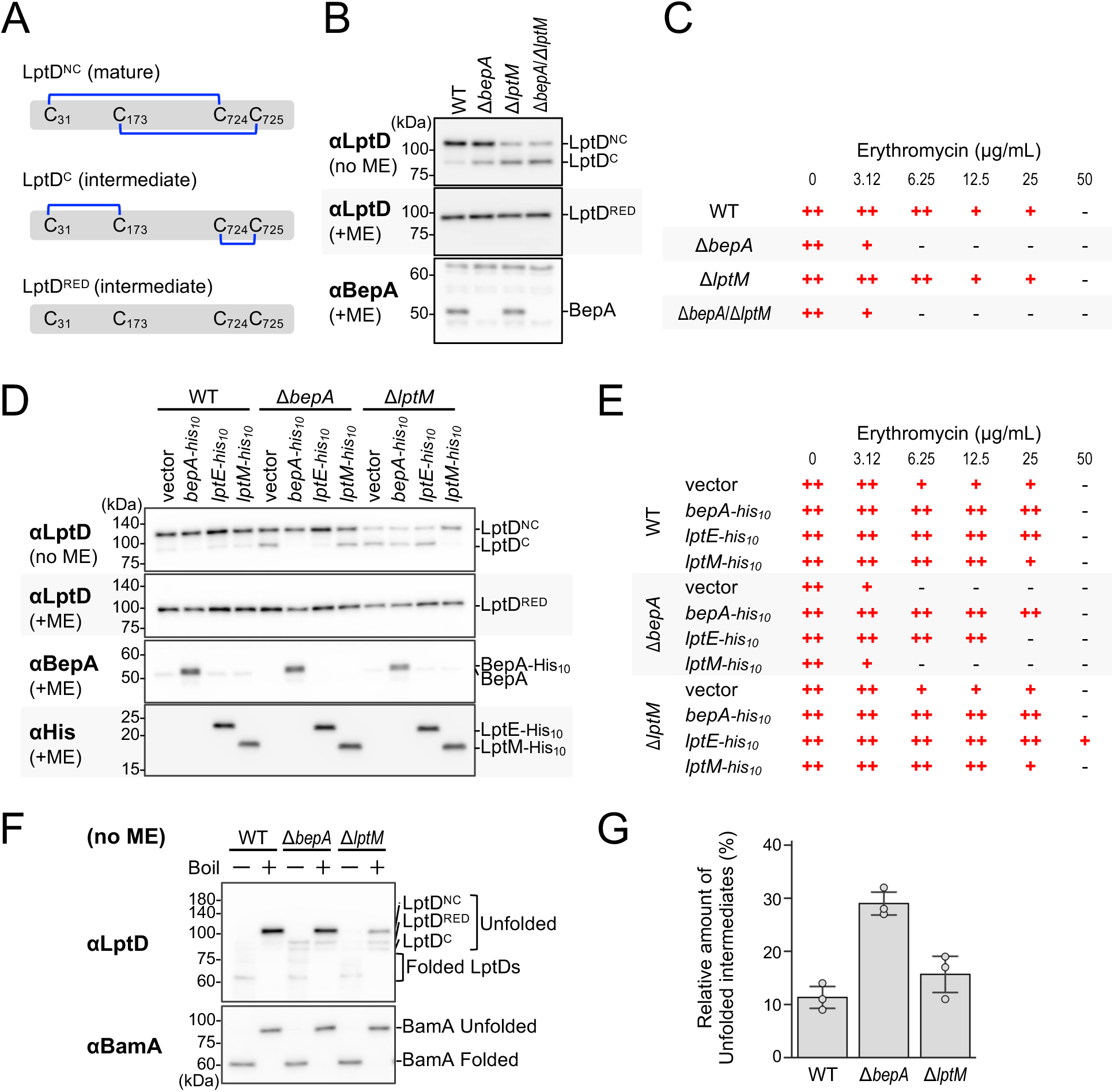
LptM and BepA act on LptD intermediates at different steps in their maturation. (**A**) Schematic representation of LptD^NC^, LptD^C^, and LptD^RED^. (**B**) Accumulation of LptD^C^ in Δ*bepA* and Δ*lptM* mutant cells. Cells of AD16 derivatives were grown at 30 °C in LB medium for 2 h. Total cellular proteins were acid-precipitated, boiled for 5 min, and then analyzed via 7.5% Laemmli SDS–PAGE under reducing (+ME) or nonreducing (no ME) conditions and immunoblotting with the indicated antibodies. (**C**) Erythromycin sensitivities of Δ*bepA* and Δ*lptM* mutant cells. Overnight cultures of AD16 derivatives were grown on LB agar plates supplemented with the indicated concentration of erythromycin, as described in the Methods section. −, no growth; +, intermediate growth; ++, wild-type growth. The original data are shown in Figure S2A. (**D, E**) Effects of BepA, LptE, or LptM overexpression on Δ*bepA* and Δ*lptM* mutant cells. (**D**) Cells of AD16 derivatives carrying pSTD689, pSTD689-*bepA-his_10_*, pSTD689-*lptE-his_10_*, or pSTD689-*lptM-his_10_* were grown at 30 °C in LB medium until the early log phase and induced with 1 mM IPTG for 1 h. Total cellular proteins were acid-precipitated and analyzed via SDS–PAGE and immunoblotting as described in **B**. (**E**) Erythromycin sensitivities of the same cells in **C** were analyzed by growing their overnight cultures on LB agar plates supplemented with 1 mM IPTG and the indicated concentration of erythromycin as in **C**. The original data are shown in Figure S2B. (**F**) Heat modifiability of the LptD intermediate in Δ*bepA* and Δ*lptM* mutant cells on SDS‒PAGE. Cells of AD16 derivatives were grown at 30 °C in LB medium for 2 h, lysed, and either boiled (denatured) or incubated at room temperature (nondenatured) for 10 min. The samples were analyzed via 7.5% Laemmli SDS–PAGE under nonreducing (no ME) conditions at 4 °C and immunoblotting with the indicated antibodies. (**G**) Relative amounts of unfolded LptD intermediates (%) were calculated according to the following equation: relative amounts of unfolded intermediates (%) = (the band intensities of unfolded LptD^RED^ and LptD^C^ in unboiled samples)/(the band intensities of unfolded LptD^NC^, LptD^RED^, and LptD^C^ in boiled samples). The mean values were plotted (error bars are S.D.; (n = 3)).

BepA is a periplasmic protease crucial for maintaining OM integrity (Narita *et al*, 2013). The overexpression of LptE complements the phenotype of Δ*bepA* cells, suggesting a functional relationship between BepA and LptD/E (Narita *et al*, 2013). BepA also interacts with the BAM complex (Narita *et al*, 2013; Daimon *et al*, 2017). Further functional and structural studies proposed a model in which BepA performs quality control of LptD during its maturation (Shahrizal *et al*, 2019; Daimon *et al*, 2020; Bryant *et al*, 2021). BepA interacts with the unstructured LptD intermediate assembled on the BAM complex. Like a chaperone, BepA facilitates the proper maturation of the LptD intermediate in the normal assembly pathway. Conversely, BepA degrades LptD intermediates abnormally stalled on the BAM complex (Miyazaki *et al*, 2021). The LptD intermediate that passes quality control by BepA interacts with LptE to form a mature LptD/E complex.

A recent study demonstrated that the small, 67-residue OM lipoprotein LptM (known as YifL before its function was understood) plays a crucial role in the LptD maturation process, interacting with LptD/E (Yang *et al*, 2023). Deletion of the *lptM* gene resulted in the abnormal accumulation of LptD^C^ and caused a synthetic phenotype in combination with deletions of either the *dsbA* gene or the *dsbC* gene, which encode a periplasmic disulfide bond oxidoreductase and an isomerase, respectively. Additionally, LptM physically and genetically interacts with the BAM complex (Yang *et al*, 2023). These findings suggest that LptM facilitates the disulfide isomerization of LptD near the BAM complex. Despite the recent structural elucidation of the *Pseudomonas aeruginosa* (*Pa*) LptD/E and *E. coli* (*Ec*) LptM heterocomplex (Luo *et al*, 2022), the precise mechanism by which LptM facilitates LptD maturation remains elusive.

Here, we investigated the timing and molecular basis of the LptM–LptD interaction. Through comprehensive mutational analyses, we defined the stage at which this interaction occurs and identified an essential region required for LptM function. By subsequently determining the cryo-EM structure of the *Ec*LptD/E/M complex and performing additional experiments, we further clarified the molecular basis of the LptM–LptD interaction and its functional role in LptD maturation. Based on these findings, we propose a model for LptM-mediated LptD assembly.

## Results and Discussion

### ① LptM acts at the late step of LptD maturation

LptM is involved in the disulfide isomerization of LptD during its maturation (Yang *et al*, 2023). In the Δ*lptM* cells, the LptD intermediate (LptD^C^) accumulated at higher levels than it did in the WT cells (Figure 1A, B). LptD^C^ accumulation was also observed in Δ*bepA* cells, albeit to a lesser extent (Figure 1A, B), as BepA interacts with LptD^C^ assembled on the BAM complex for quality control (Miyazaki *et al*, 2021). Notably, the Δ*lptM* cells showed erythromycin sensitivity similar to WT cells, whereas the Δ*bepA* cells exhibited higher sensitivity (Figure 1C). Erythromycin is a macrolide antibiotic commonly used to assess OM integrity. These findings implied that the accumulated LptD^C^ in Δ*lptM* and Δ*bepA* cells, detected via SDS‒PAGE, may include forms derived from distinct intermediate states of LptD. Thus, LptM and BepA would independently act at different steps of LptD maturation. In support of this hypothesis, Δ*lptM* and Δ*bepA* double mutant cells did not show additive or synergistic effects (Figure 1B, C).

We further investigated the independence of LptM and BepA functions in LptD maturation. In Δ*lptM* cells, the overproduction of LptM-His_10_ suppressed LptD^C^ accumulation, whereas that of BepA-His_10_ had no effect (Figure 1D). In contrast, both LptD^C^ accumulation and erythromycin sensitivity in Δ*bepA* cells were suppressed by the overproduction of BepA-His_10_ but not by that of LptM-His_10_ (Figure 1D, E). In addition, LptE overproduction indeed suppressed both LptD^C^ accumulation and erythromycin sensitivity in Δ*bepA* cells (Figure 1D, E), as reported previously (Narita *et al*, 2013), whereas it did not suppress LptD^C^ accumulation in Δ*lptM* cells (Figure 1D). These results support the hypothesis that LptM and BepA act independently during LptD maturation.

Next, we investigated the stage at which LptM acts during LptD maturation. β-barrel OMPs solubilized with a mild detergent retain compact barrel shapes, allowing them to migrate faster on SDS‒ PAGE than the same samples subjected to heat denaturation by boiling (Noinaj *et al*, 2015). Using this property, we performed a heat modifiability assay to analyze the structural states of the LptD intermediate in each mutant strain (Figure 1F, Figure S3). As a control, BamA exhibited faster mobility in all cells when the samples were not boiled, confirming that it forms a β-barrel structure (a compact folded structure) even on SDS‒PAGE. In WT cells, mature LptD (LptD^NC^) also formed a compact folded structure with faster SDS‒PAGE mobility when the samples were not boiled under nonreducing conditions (Figure 1F). In both Δ*bepA* and Δ*lptM* cells, heat-denaturing unfolded LptD^NC^ and LptD intermediates (LptD^RED^ and LptD^C^) were detected when the samples were boiled. Notably, the amount of LptD^RED^ and LptD^C^ in Δ*bepA* cells on SDS‒PAGE remained unchanged regardless of whether the samples were boiled (Figure 1F). This result indicates that most accumulated LptD intermediates in Δ*bepA* cells were not in folded forms, which is consistent with previous studies showing that BepA interacts with unstructured LptD intermediates assembled on the BAM complex (Miyazaki *et al*, 2021). In contrast, the unfolded LptD intermediates were hardly detected in the Δ*lptM* cells as they were in the WT cells when the samples were not boiled (Figure 1F). Quantifying the percentage of unfolded LptD intermediates to total LptD in the cells revealed that the rate was greater in the Δ*bepA* cells than in the WT cells, whereas it was nearly identical between the Δ*lptM* cells and WT cells (Figure 1G), suggesting that LptD intermediates in folded forms predominantly accumulated in the Δ*lptM* cells. Taken together, these results imply that LptM acts on β-barrel LptD intermediates at a late step of LptD maturation, following the action of BepA.

### ② An N-terminal conserved region of LptM is essential for its function

LptM is a small OM lipoprotein composed of fewer than 50 amino acid residues, excluding its signal sequence (Figure 2A). To gain insight into how this small lipoprotein facilitates LptD maturation, we constructed C-terminal systematically truncated mutants of LptM by introducing a stop codon, an *amber* codon, into its ORF and examined their function (Figure 2B). Most truncated LptM mutants suppressed the abnormal accumulation of LptD^C^ in Δ*lptM* cells, similar to WT LptM and LptM-His_10_. Interestingly, even the LptM P29*amber* mutant, which contains only nine amino acids (C_20_GLKGPLYF_28_), maintained its function (Figure 2B). Importantly, this N-terminal region of LptM is highly conserved (Figure 2A) (Yang *et al*, 2023). Hence, the conserved N-terminal short region is essential for LptM function. To elucidate the molecular basis of LptM-mediated LptD maturation, structural information on the *Ec*LptD/E/M complex is essential.

**Figure 2.**
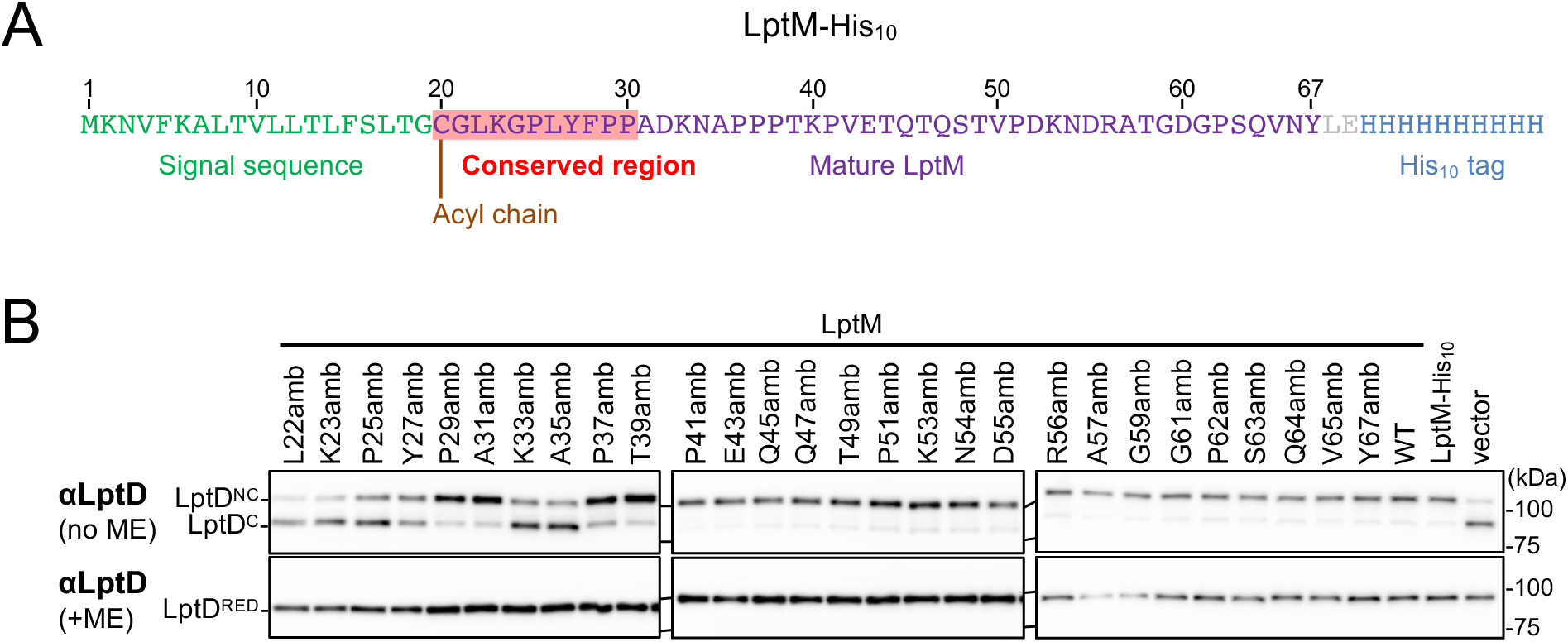
Essential region for LptM function. (**A**) Amino acid sequence of *E. coli* LptM-His_10_. The signal sequence and mature region are colored green and purple, respectively. The red marker indicates the highly conserved region of LptM. The Cys-20 residue at the N-terminus of mature LptM is a triacylated site. (**B**) Activities of LptM truncated mutants. The RM4749 (Δ*lptM*) cells carrying the pTWV228 or pTWV228*-lptM(amb)-his_10_* plasmids were grown at 30 °C in LB medium until the early log phase and induced with 1 mM IPTG for 1 h to express the indicated LptM mutants. Total cellular proteins were acid-precipitated and analyzed by 7.5% Laemmli SDS-PAGE under reducing (+ME) or nonreducing (no ME) conditions and immunoblotting with anti-LptD antibodies.

### ③ Cryo-EM structure of the LptD/E/M complex

Previous structural studies of the LptD/E complex revealed that LptD consists of a periplasmic β-taco domain at its N-terminus and a 26-stranded β-barrel transmembrane domain at its C-terminus, which are connected by two intramolecular disulfide bonds, with LptE embedded within the β-barrel domain of LptD (Figure S4A) (Dong *et al*, 2014; Qiao *et al*, 2014; Botos *et al*, 2016). The crystal structure of the *Pa*LptDE complex in association with *Ec*LptM revealed that LptM is positioned at the interface between the β-taco and β-barrel of LptD (Luo *et al*, 2022). However, information from interspecies complexes remains limited, particularly regarding their relevance and physiological significance, which have yet to be fully elucidated. To address this, we determined the cryo-EM structure of the *Ec*LptD/E/M complex at 3.27 Å resolution (Figure 3A).

**Figure 3.**
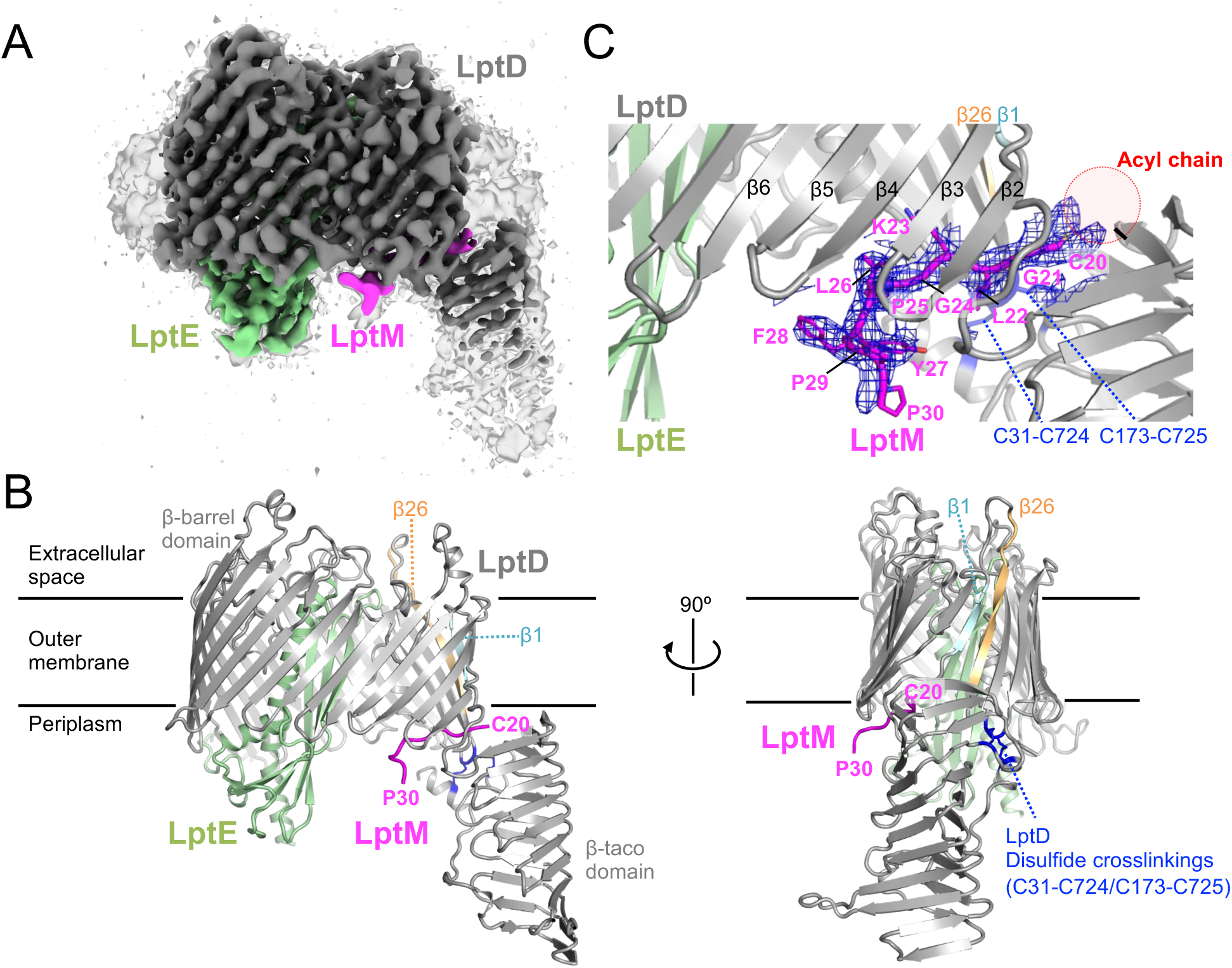
Cryo-EM structure of the *E. coli* LptD/E/M complex. (**A**) Cryo-EM map of the nanodisc-reconstituted *Ec*LptD/E/M complex. (**B**) Cartoon model of *Ec*LptD/E/M. LptD, LptE, and LptM are shown in gray, light green, and magenta, respectively. The 1st β-strand (β1) and final β-strand (β26) of the β-barrel domain of LptD and two intramolecular disulfide bonds in LptD are shown in cyan, orange, and blue, respectively. (**C**) Enlarged view of the LptD/LptM interface. The side chains of the amino acid residues of LptM on the interface between LptD and LptM are shown as stick models with the cryo-EM map (blue mesh, contoured at 5.0σ). A putative position of the acyl chains on the N-terminus of LptM is shown in a red circle.

Our cryo-EM structure revealed that the LptD/E complex adopts a conformation consistent with previously reported structures (Dong *et al*, 2014; Qiao *et al*, 2014; Botos *et al*, 2016), with LptM positioned between the two domains of LptD (Figure 3B). Notably, only the N-terminal conserved, essential region (C20–P30) of LptM was observed: the ordered feature of this conserved region suggests that the LptD/E/M structure provides critical insights into the functional role of LptM. The side chain of Cys-20, located at the N-terminus of LptM, is oriented toward the outer membrane (OM) layer, outside the β-barrel domain of LptD. It is reasonable that Cys-20 is acylated and anchored to the OM at this position. Indeed, the EM map shows a region corresponding to the acylated site, although it is not clearly defined (Figure 3C). In the crystal structure of *Pa*LptD/E-*Ec*LptM, the ordered triacyl chain was found at the same β-barrel/β-taco interface (Luo *et al*, 2022). The LptM essential region interacts with the N-terminal β-strands (β2–4) of the LptD β-barrel domain and is located near the β-barrel seam (β1/β26) and the periplasmic taco domain containing intramolecular disulfide bonds in LptD (Figure 3B, C). These regions of LptD are highly conserved among LptM orthologs (Botos *et al*, 2016). The positioning of LptM was supported by *in vivo* photo-crosslinking between LptM L22*p*BPA and LptD (Figure S5) (Yang *et al*, 2023). This proximity to the disulfide bonds (C31-C724, C173-C725) of LptD implies that LptM contributes to the maturation of LptD, including the disulfide isomerization of LptD.

As shown in Fig. 1, LptM acts at the late stage of LptD maturation, possibly contributing to its β-barrel formation. Here, we focused on the junction between β1 and β26 of LptD, which is likely formed at the final stage of β-barrel assembly. Comparison of our *Ec*LptD/E/M cryo-EM structure with the crystal structure of *Ec*LptD(Qiao *et al*, 2014) and the AlphaFold2 model of *Ec*LptD/E/M (Yang *et al*, 2023) revealed that the β-barrel junction in the cryo-EM structure was the most tightly closed (Figure S4A, B). This finding suggests that LptM strengthens and stabilizes the folding of LptD by acting as a "barrel-rivet". Consistent with this finding, recent hydrogen–deuterium exchange mass spectrometry analysis using purified *Ec*LptD/E/M and *Ec*LptD/E suggested that LptM stabilizes the β-barrel/β-taco regions in proximity to the lateral gate of LptD (Yang *et al*, 2023). A comparison of the *Pa*LptD/E-*Ec*LptM and apo *Pa*LptD/E structures also revealed the dynamic nature of the β-taco domain (Luo *et al*, 2022).

The nonessential C-terminal region (from residue 31 to the C-terminus) is disordered in the cryo-EM structure, and the functional role of this flexible region remains unclear. In an AlphaFold2 model of the *Ec*LptD/E/M complex, the less conserved C-terminal region extends into the β-barrel domain inside LptD, which is supported by *in vivo* photo-crosslinking (Yang *et al*, 2023). This positioning implies that the C-terminal region of LptM assists in LptD maturation and LPS transport through the LptD/E complex. In addition, several residues in the C-terminal region of LptM are crosslinked to OmpA (Yang *et al*, 2023), suggesting that part of this region can be temporarily exposed outside the LptD β-barrel. This idea is reasonable because of the flexibility of the C-terminal region. Further study is needed to understand the contribution of the C-terminal region to LptD maturation and LPS transport.

### ④ Interaction between LptD and LptM facilitates LptD maturation

To further verify whether *Ec*LptD/E/M forms in the EM structure in living cells, we performed *in vivo* disulfide crosslinking experiments using Cys derivatives of LptM and LptD. A Cys mutation was introduced at F28 in the essential region of LptM-3xFLAG, and six residues (Y248, P253, L279, Q366, N443, and S617) on the periplasmic-facing side of the β-barrel domain in LptD-His_10_ (Figure 4A). These LptM and LptD derivatives were functional, as they complemented the phenotypes of Δ*lptM* and *lptD*- depleted cells, respectively (Figure S6A, B). We coexpressed a combination of each LptM-3xFLAG and LptD-His_10_ Cys derivative in Δ*lptM*, *lptD*-depleted cells and examined disulfide bond formation between LptM and LptD. Under nonreducing conditions, higher-mass products reactive to both anti-FLAG and anti-His antibodies were detected in two combinations: LptM(F28C)/LptD(P253C) and LptM(F28C)/LptD(L279C) (Figure 4B). These bands disappeared upon 2-mercaptoethanol (ME) treatment, confirming that they were disulfide crosslinked products of LptM-3xFLAG and LptD-His_10_. These *in vivo* results are consistent with the EM structure in which F28 of LptM is close to P253 and L279 of LptD (Figure 4A).

**Figure 4.**
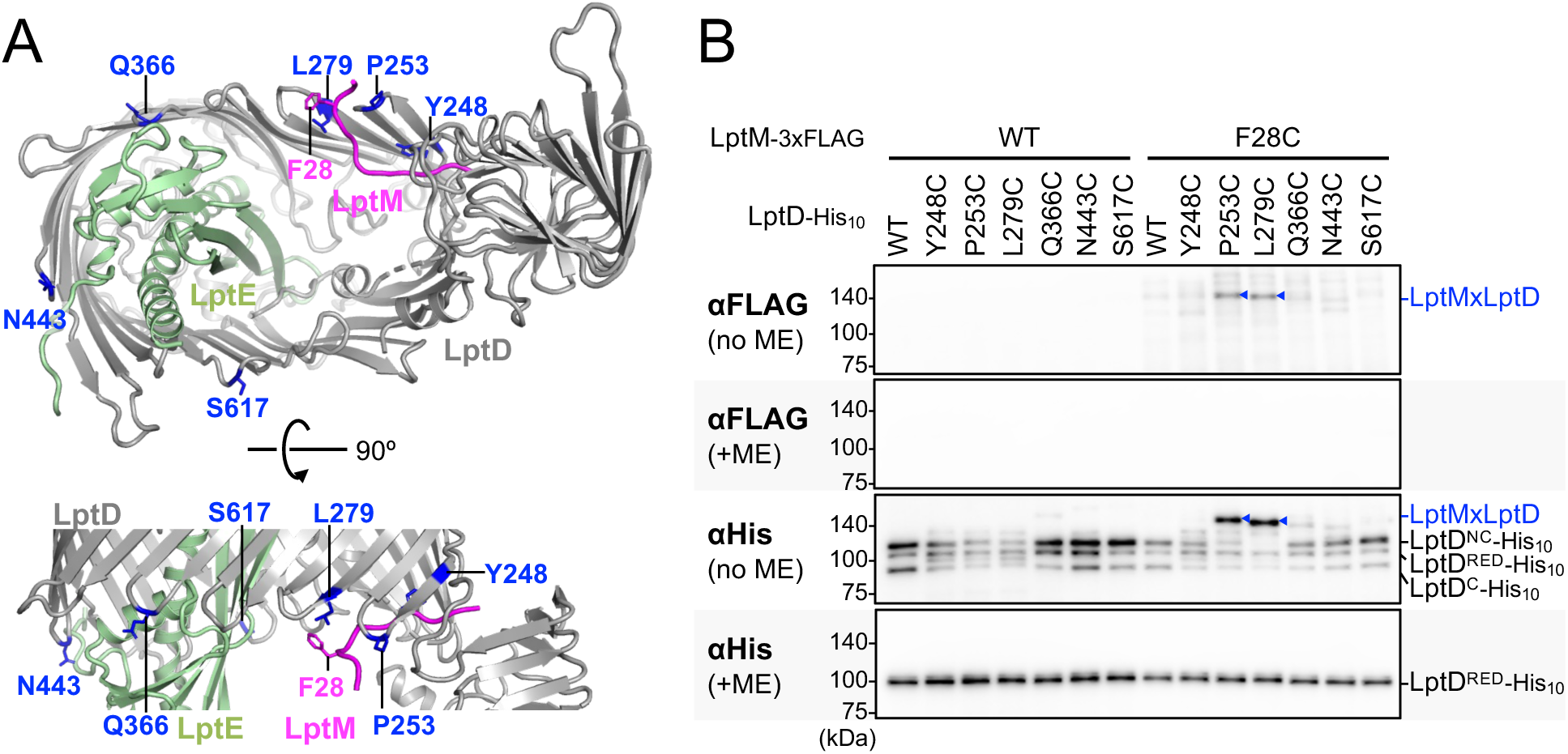
The conserved LptM region interacts with the N-terminal regions of the LptD β-barrel domain. (**A**) Mapping of the Cys-introduced positions on LptM and LptD in the *E. coli* LptD/E/M structure. LptD, LptE, and LptM are shown in gray, light green, and magenta, respectively. The side chains of Phe-28 of LptM are shown as stick models. Cys-introduced sites on LptD are shown in blue, with the side chains used as stick models. (**B**) Disulfide crosslinking between the Cys residues in the essential region of LptM and the β-barrel domain of LptD. Cells of RM5205 (P*_ara_-lptD*, Δ*lptM*) carrying a combination of plasmids encoding WT or a Cys-introduced mutant of LptM-3xFLAG and LptD-His_10_ as indicated were grown at 30 °C in LB medium and induced with 1 mM IPTG for 3 h to express LptM(Cys)-3xFLAG and LptD(Cys)-His_10_. Total cellular proteins were acid-precipitated, solubilized with SDS-buffer containing NEM (for blocking free thiol groups), treated with or without 2-mercaptoethanol (ME), analyzed by 7.5% Laemmli SDS‒PAGE under reducing (+ME) or nonreducing (no ME) conditions and immunoblotting with the indicated antibodies.

Furthermore, to investigate the significance of the LptM–LptD interaction, we examined the function of LptM by introducing Ala, Cys, or Trp mutations in its essential region. Ala, Cys, and Trp mutations, with the exception of G21W, had only minor effects on LptM function, with no significant effect observed (Figure 5A–C). The G21W mutation significantly impaired LptM activity (Figure 5C). In addition, introducing this G21W mutation into LptM L22*p*BPA reduced its crosslinking efficiency with LptD. In contrast, the G21W mutation did not affect the crosslinking of LptM K53*p*BPA with OmpA, an OMP (Figure 5D). These findings suggest that the G21W mutation specifically disrupts the LptM–LptD interaction but does not affect the OM localization of LptM. Taken together, these findings indicate that the essential region of LptM interacts with LptD to facilitate its proper assembly and maturation.

**Figure 5.**
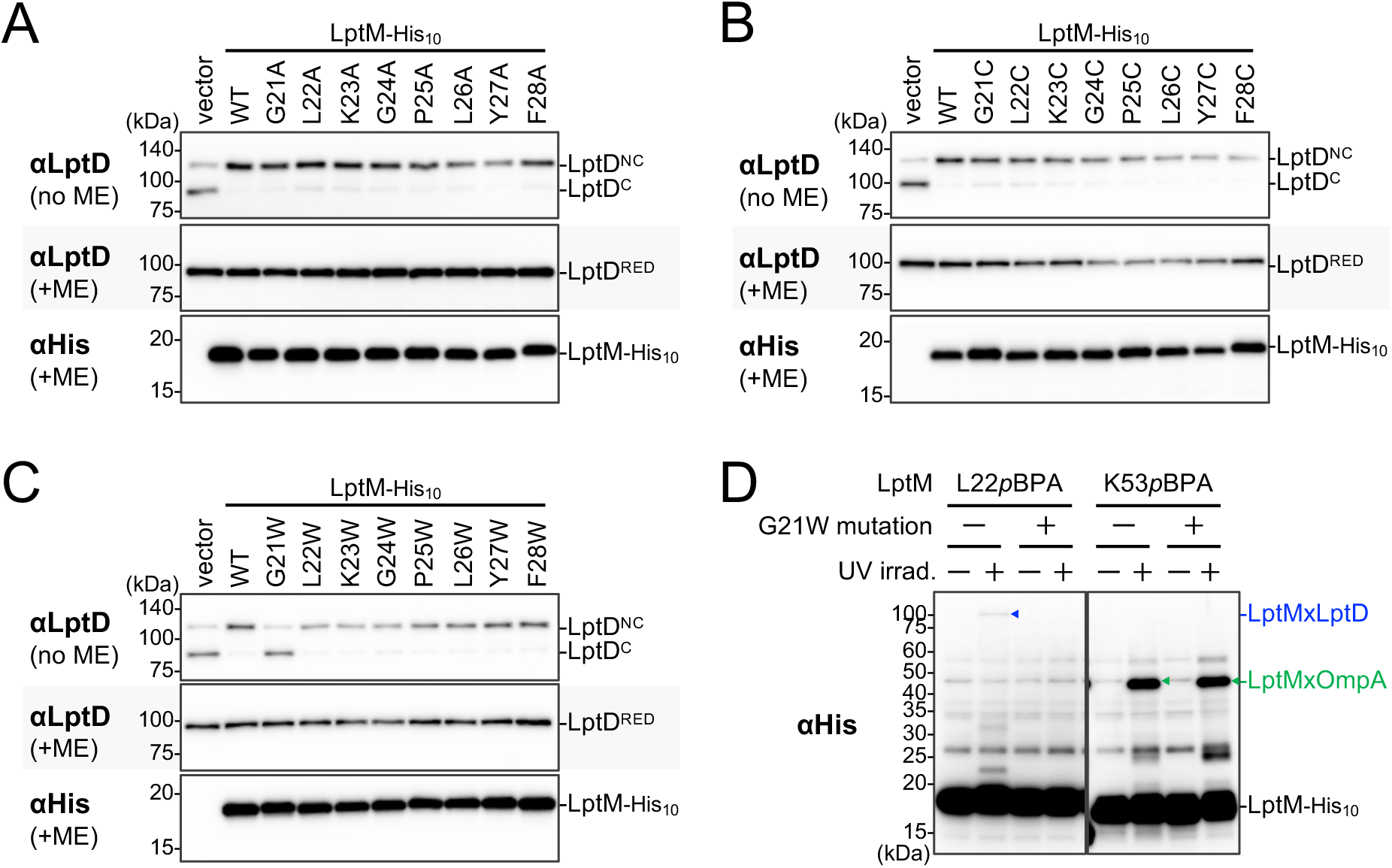
The conserved region of LptM is crucial for its function and interaction with LptD. (**A–C**) Systematic mutational analysis of the conserved region of LptM. Cells of RM4749 (Δ*lptM*) carrying pTWV28, pTWV28*-lptM(Ala)-his_10_*, pTWV28*-lptM(Cys)-his_10_* or pTWV28*-lptM(Trp)-his_10_* plasmids were grown and analyzed via SDS–PAGE and immunoblotting, as shown in Figure 2B. (**D**) Effect of a G21W mutation on LptM photo-crosslinking. The RM4749 (Δ*lptM*) cells harboring pEVOL-pBpF and pTWV28-*lptM(mut, amb)-his_10_* plasmids were grown at 30 °C in LB medium supplemented with 0.02% arabinose and 0.5 mM *p*BPA until the early log phase was reached, after which they were induced with 1 mM IPTG for 1 h to express the indicated LptM(*p*BPA) variants. The cultures were divided into two portions, each of which was treated with or without UV irradiation for 10 min at 4 °C. Total cellular proteins were acid-precipitated and analyzed by 7.5% Laemmli SDS-PAGE and immunoblotting with anti-His antibodies.

### ⑤ Model for the proposed LptD assembly process

As LptD is an essential OMP and a promising target of novel antibiotics, its maturation process has been well studied (Lee *et al*, 2016, 2018, 2019). These studies have provided important insights into the BAM-mediated assembly of OMPs into the OM. Based on these findings, we propose a model for the LptD assembly process (Figure S7). The precursor of LptD is synthesized in the cytoplasm and delivered to the SecYEG translocon. After its N-terminal signal sequence is cleaved off, LptD is translocated through SecYEG to the periplasm, where DsbA oxidizes the Cys residues of LptD^RED^ to form LptD^C^ (Ruiz *et al*, 2010). LptD is then delivered to the BAM complex by periplasmic chaperones, including SurA (Figure S7 (i)) (Schwalm *et al*, 2013; Fenn *et al*, 2024). Like other OMPs, the C-terminal region of the β-barrel region of LptD directly interacts with the BAM complex (Lee *et al*, 2016, 2019; Miyazaki *et al*, 2021). At this stage, BepA interacts with the C-terminal half of the β-barrel domain of LptD assembled on the BAM complex. BepA either facilitates the maturation of LptD when its assembly normally proceeds or degrades misassembled LptD, which abnormally remains stalled on BAM (Figure S7 (ii)) (Miyazaki *et al*, 2021). After discrimination by BepA, the N-terminal region of the LptD β-barrel undergoes further maturation within the β-barrel domain of BamA. Finally, the LptD intermediate on BAM encounters its partner protein LptE to form a complex (Figure S7 (iii)) (Lee *et al*, 2016, 2019).

During LptD maturation, while LptD interacts with LptE, which facilitates the disulfide isomerization of LptD^C^ to LptD^NC^, LptM functions as a crucial factor (Chng *et al*, 2012; Yang *et al*, 2023). Our results suggest that LptM acts on the folded LptD intermediate after BepA-mediated quality control (Figure 1). Furthermore, the LptD/E complex partially remains associated with the BAM complex through interactions with BamB and BamD (Yang *et al*, 2023). Importantly, unlike Δ*bepA* or *lptD4213* mutant cells, in which stalled LptD intermediates disrupt BAM function and increase erythromycin sensitivity, Δ*lptM* cells do not present such phenotypes (Figure 1). These findings suggest that LptD intermediates in Δ*lptM* cells do not occupy the BAM complex, thereby allowing the normal assembly of other nascent OMPs. We propose that LptM interacts with the LptD/E complex, forming a mature structure near BAM, to facilitate the disulfide isomerization of LptD^C^ to LptD^NC^ (Figure S7 (iv), (v)). Considering that this short essential region tightly interacts with the β-barrel domain of LptD (Figure 3, 4), it is unlikely to serve as a recruiter for the disulfide oxidase DsbA or disulfide isomerase DsbC to the LptD intermediate. Although no amino acid residues of the essential LptM region, except for G21, have a decisive impact (Figure 5), its characteristic sequence broadly recognizes LptD through both its main chain and side chains. This raises the question of how LptM facilitates the disulfide isomerization of LptD. In our LptD/E/M structure, the β- barrel domain of LptD is more tightly closed than the LptD/E complex without LptM. Thus, LptM may structurally stabilize the LptD/E conformation, which is suitable for disulfide isomerization.

The C-terminal β-strand of the substrate OMP interacts with the N-terminal β1 of BamA in the first step of the BAM-mediated OMP assembly process (Doyle *et al*, 2022; Shen *et al*, 2023). Following maturation, the substrate OMP is released from BAM into the OM, where its separated N-terminal/C-terminal β-strands associate to form a closed β-barrel structure. Given that LptM directly interacts with the N-terminal β-strands in the LptD β-barrel (Figures 3–4), the LptM–LptD interaction may assist in the β- barrel closure of LptD and its release from BAM, ensuring the proper assembly and function of LptD.

In this study, we elucidated key aspects of the mechanism underlying LptM-mediated LptD maturation. Notably, the conserved N-terminal region of LptM is sufficient to chaperone LptD maturation. This discovery provides valuable insights into the development of novel OM-targeting antibiotics against the LptD/E/M complex, analogous to darobactin, which inhibits the BAM complex (Kaur *et al*, 2021). Moreover, our findings suggest the intriguing possibility that unidentified very short lipoproteins or secreted proteins, consisting almost entirely of signal sequences, may play critical roles in the maturation of larger proteins, such as LptM, or in other essential cellular processes. Investigating these microproteins represents an exciting avenue for future research.

## Methods

### Bacterial strains, Plasmids and Primers

The *Escherichia coli* K12 strains, plasmids, and primers used in this study are listed in Tables S1, S2, and S3, respectively. Details of the strain and plasmid construction and media are described in the Construction of Mutant Strains and Plasmid Construction sections, respectively.

### Media and Bacterial Cultures

*E. coli* cells were grown in LB medium (Nacalai Tesque). 50 or 100 µg/mL ampicillin (Amp), 20 µg/mL chloramphenicol (Cm), 25 µg/mL kanamycin (Km), 25 µg/mL tetracycline (Tet), and/or 50 µg/mL spectinomycin (Spc) were added as appropriate for growing plasmid-bearing cells and selecting transformants and transductants. Bacterial growth was monitored with a Mini photo 518R (660 nm; TAITEC Co.).

### Antibodies

The anti-His-tag pAb (PM032) was purchased from MBL, and the anti-FLAG M2 antibody was purchased from Sigma‒Aldrich. The anti-BepA (Narita *et al*, 2013), anti-LptD (Narita *et al*, 2013), and anti-BamA (Gunasinghe *et al*, 2018) antibodies were used as described previously.

### Construction of Mutant Strains

RM4717 (DY330, Δ*lptM*::*kan*) was constructed via essentially the same procedure as the construction of strains with a chromosomal C-terminal His_10_-tagged gene (Miyazaki *et al*, 2020b). First, a Δ*lptM*::*kan* fragment, which has a sequence identical to the upstream of the *lptM* start codon and the downstream regions of the *lptM* stop codon at the respective ends of the fragment, was PCR-amplified from pKD4 (a plasmid carrying a *kan* cassette) (Datsenko & Wanner, 2000)using a pair of primers, d-lptM-4F and d-lptM- 4R. The chromosomal *lptM* locus of the *E. coli* DY330 strain was subsequently replaced with this fragment via the λ-Red recombination system (Yu *et al*, 2000). RM4749 (AD16, Δ*lptM*::*kan*) and RM4752 (AD16, Δ*bepA* Δ*lptM*::*kan*) were constructed by transducing Δ*lptM*::*kan* from RM4717 to AD16 (Kihara *et al*, 1995) and SN56 (Narita *et al*, 2013), respectively. RM5181 (HM1742, *araC-*P*_araBAD_-lptD*) was constructed by removing the *kan* cassette of RM3588 (HM1742, *kan araC*-P*_araBAD_-lptD*) (Miyazaki *et al*, 2021) using pCP20 (Cherepanov & Wackernagel, 1995). RM5205 (HM1742, *araC-*P*_araBAD_-lptD* Δ*lptM*::*kan*) was constructed by transducing Δ*lptM*::*kan* from RM4717 to RM5181.

### Plasmid Construction

pRM1441 (pTWV228-*lptM-his_10_*) and pTWV228-*lptM(amb)-his_10_* plasmids (pRM1442–pRM1470) were synthesized by GenScript. pTWV228-*lptM(Ala)-his_10_* (pRM1531–pRM1538), pTWV228-*lptM(Cys)-his_10_* (pRM1539–pRM1546), and pTWV228-*lptM(Trp)-his_10_* (pRM1547–pRM1554) were constructed from pRM1441 via site-directed mutagenesis. pRM1601 (pTWV228-*lptD(G21W, L22amb)-his_10_*) and pRM1604 (pTWV228-*lptD(G21W, K53amb)-his_10_*) were constructed from pRM1550 (pTWV228-*lptD(G21W)-his_10_*) via site-directed mutagenesis.

Derivatives of pRM267 (pTWV228-*lptD-his_10_*) (Daimon *et al*, 2017) carrying an additional Cys mutation introduced into the *lptD* gene (pRM1577, pRM1588, and pRM1580–pRM1583) were constructed via site-directed mutagenesis.

pRM867 (pSTD689-*bepA-his_10_*) was constructed by subcloning the EcoRI-SalI *bepA-his_10_* from pUC-bepA-his_10_ (Narita *et al*, 2013) into the same site of pSTD689 (Kanehara *et al*, 2003). pRM1381 (pSTD689-*lptE-his_10_*) was constructed as follows. A pSTD689-based vector fragment was PCR-amplified from pRM868 (pSTD689-*bepA(E137Q)-his_10_*) using a pair of primers, pSTD-H-F and pSTD-H-R. An *lptE* fragment was prepared via PCR amplification from the genome of MC4100 using a pair of primers, lptE-F and lptE-R. These two fragments were ligated via *in vitro* recombination using an In-Fusion HD cloning kit (Takara Bio) to obtain pRM1381. pRM1383 (pSTD689-*lptM-his_10_*) was also constructed by *in vitro* recombination of the above pSTD689-based vector fragment and an *lptM* fragment prepared by PCR amplification from the genome of MC4100 using a pair of primers, lptM-F and lptM-R. To obtain pRM1555 (pSTD689-*lptM-3xflag*), the pSTD689-*lptM* sequence with a *3xflag* tag at the C-terminus of *lptM* was PCR amplified from pRM1383 using a pair of primers, lptM-3F-F and lptM-3F-R, self-ligated via *in vitro* recombination using an In-Fusion HD cloning kit. pRM1566 (pSTD689-*lptM(F28C)-3xflag*) was constructed from pRM1555 via site-directed mutagenesis.

pRM1606 (pTV118N-*lptD/lptE/lptM-his_10_*; Figure S8A) was constructed as follows. A vector fragment was prepared via PCR amplification from pTV118N (Takara Bio) using a pair of primers, pTV-F and pTV-R. *lptD*, *lptE*, and *lptM-his_10_* fragments were amplified via PCR from pRM309 (a plasmid encoding *lptD*; (Miyazaki *et al*, 2018)), pRM1381, and pRM1383, respectively, using pairs of primers, namely, lptD-F/lptD-R, lptE-F2/lptE-R2, and lptM-F2/lptE-R2. These four fragments were ligated via *in vitro* recombination using an In-Fusion HD cloning kit; however, we obtained only a plasmid lacking *lptE* (pRM1557). A vector fragment containing the *lptD* and *lptM-his*_10_ genes and an *lptE* fragment were amplified via PCR from pRM1557 and pRM1381 using pairs of primers, namely, plptDM-F/plptDM-R and lptE-F3/lptE-R3, respectively. These were ligated via *in vitro* recombination using an In-Fusion HD cloning kit to obtain a plasmid encoding *lptD*, *lptE*, and *lptM-his_10_*. After sequential site-direct mutagenesis for the correction of unexpected mutations, we obtained pRM1606.

### Immunoblotting analysis

Acid-denatured proteins, whose preparation details for each experiment are shown in the figure legends, were solubilized in SDS-sample buffer (62.5 mM Tris-HCl (pH 6.8), 2% SDS, 10% glycerol, and 5 mg/mL bromophenol blue) with or without 10% β-ME, boiled at 98 °C for 5 min, separated by SDS‒PAGE and electroblotted onto a PVDF membrane (Merck Millipore). The membrane was first blocked with 1% skim milk in PBST (phosphate-buffered saline with Tween 20) and then incubated with anti-His-tag pAb (PM032) (1/25,000 dilution), anti-FLAG M2 antibody (1/100,000), anti-BepA (1/10,000), anti-LptD (1/50,000 or 1/100,000) or anti-BamA (1/100,000). After being washed with PBST, the membrane was incubated with a horseradish peroxidase (HRP)-conjugated secondary antibody (1/5,000) (goat anti-rabbit IgG (H + L)-HRP conjugate; Bio-Rad) in PBST. After washing with PBST, the proteins were visualized with detection reagents, Chemi-Lumi One (Nacalai Tesque), and a chemiluminescence image analyzer, FUSION Solo S (VILBER).

### Purification and nanodisc reconstitution of the LptD/E/M complex

The workflow for purification and nanodisc reconstitution of the EcLptD/E/M complex is shown in Figure S8B. The cells of KRX (Promega) carrying pRM1606 (Figure S8A) were inoculated into LB medium containing 0.4% glucose and 50 μg/mL ampicillin and cultivated at 37 °C for 16 h. The overnight culture was inoculated with LB medium containing 50 μg/mL ampicillin and grown at 30 °C until the OD_600_ reached 1.0. Isopropyl β-D-thiogalactopyranoside (IPTG) was then added at a concentration of 1 mM, and the cells were further cultured at 30 °C for 3 h to induce the expression of LptD, LptE, and LptM-His_10_. The cells were collected via centrifugation, suspended in buffer (20 mM Tris-HCl (pH 8.0), 150 mM NaCl, 1 mM EDTA-Na (pH 8.0), 0.1 mM phenylmethylsulfonyl fluoride (PMSF)), and disrupted with a Microfluidizer Processor M-110EH at 100 MPa (Microfluidics International). After removal of the unbroken cells and protein aggregations by centrifugation, the collected supernatant was further ultracentrifuged (40,000 rpm for 60 min, 4 °C, Beckman 45Ti rotor) to obtain the membrane fraction. The membrane fraction was resuspended in solubilization buffer (50 mM Tris-HCl (pH 8.0), 150 mM NaCl, 1% (w/v) n-dodecyl β-maltoside (DDM), 10 mM imidazole-HCl (pH 8.0), 0.1 mM PMSF) and solubilized by gentle stirring at 4 °C for 60 min. After removal of the insoluble fraction by ultracentrifugation (40,000 rpm for 30 min, 4 °C, Beckman 45Ti rotor), the supernatant was mixed with 2.5 ml of Ni-NTA agarose resin (QIAGEN) preequilibrated with solubilization buffer and gently stirred at 4 °C for 60 min. Then, 2.5 mL of wash buffer (50 mM Tris-HCl (pH 8.0), 150 mM NaCl, 0.05% (w/v) DDM, 50 mM imidazole-HCl (pH 8.0), 0.1 mM PMSF) was added to the column 3 times. Next, 2.5 mL of elution buffer (50 mM Tris-HCl (pH 8.0), 150 mM NaCl, 0.05% (w/v) DDM, 500 mM imidazole-HCl (pH 8.0), 0.1 mM PMSF) was added to the column 6 times. The eluted fractions containing LptD, LptE, and LptM-His_10_ were collected and concentrated with an Amicon Ultra 50K NMWL (Merck Millipore). After ultracentrifugation (45,000 rpm for 30 min, himac S55A2 rotor), the concentrated sample was then applied to a Superdex 200 Increase 10/300 GL column (Cytiva) equilibrated with buffer (50 mM Tris-HCl (pH 8.0), 150 mM NaCl, 0.01% (w/v) DDM, 0.1 mM PMSF). The fractions eluted with the target proteins were collected and concentrated with an Amicon Ultra 50K NMWL.

A protein‒lipid mixture containing 11.6 mg/mL purified *Ec*LptD/E/M complex, 8.7 mg/mL purified MSP1D1, a membrane scaffold protein (Nath *et al*, 2007), and 25 mg/mL *E*. *coli* phospholipids (Avanti) in buffer (50 mM Tris-HCl (pH 8.0), 150 mM NaCl, 0.01% (w/v) DDM, 0.1 mM PMSF) was gently mixed at 4 °C for 1 h. Bio-Beads SM2 (Bio-Rad) were added to the mixture to remove the detergent. After gently mixing at 4 °C overnight, the mixture was filtered through 0.22 μm PVDF centrifugal filters (Millipore) to completely remove the Bio-Beads SM2. After ultracentrifugation (45,000 rpm for 30 min, himac S55A2 rotor), the sample was loaded on a Superdex 200 10/300 GL column equilibrated with buffer (50 mM Tris-HCl (pH 8.0), 150 mM NaCl, 0.1 mM PMSF). After analysis via SDS‒PAGE and native PAGE (Figure S8C, D), the fractions containing the *Ec*LptD/E/M-reconstituted nanodiscs were collected and concentrated with an Amicon Ultra 100K NMWL (Merck Millipore).

### Cryo-EM grid preparation

Quantifoil holey carbon grids (Cu R1.2/R1.3, 300 mesh) were glow-discharged at 7 Pa with 10 mA for 10 seconds using a JEC-3000FC sputter coater (JEOL) before sample application. A 3-μL aliquot of the nanodisc-reconstituted *Ec*LptD/E/M complex (approximately 3 mg/mL) was applied onto the grids, which were blotted for 3 seconds at 100% humidity and 8 °C, blot force 10, and plunged into liquid ethane via a Vitrobot Mark IV (Thermo Fisher Scientific).

### Cryo-EM data collection and processing

The cryo-EM dataset was collected on a Glacios Cryo-TEM instrument operated at an accelerating voltage of 200 kV and equipped with a Falcon4 direct electron detector (Thermo Fisher Scientific) at the Institute for Life and Medical Sciences, Kyoto University. Images were collected at a nominal magnification of 150,000×, corresponding to a calibrated pixel size of 0.924 Å/pixel, 62 frames per image with a total exposure dose of 50 e−/Å^2^, via the EPU software (Thermo Fisher Scientific).

The data processing was performed with the cryoSPARC v4.5.3 software platform (Punjani *et al*, 2017). Additionally, the Cryo-EM data-processing workflow for the *Ec*LptD/E/M complex is shown in Figure S9. A total of 4,474 movies were aligned using patch motion correction, and the contrast transfer function (CTF) parameters were estimated using the Patch CTF estimation. From 200 of the 4,474 micrographs, 107,407 particles were automatically picked up using Blob picker in cryoSPARC. The particles were subjected to two rounds of 2D classifications, generating 2D class averages. Using these 2D class averages as templates, the particles were re-picked with the Template Picker form the 4,474 micrographs, resulting in 1,725,039 particles. The particles were then subjected to 2D classifications. The only 500,000 particles were then subjected to Ab-intio reconstruction (K=6), generating 3D initial models. The particles were then subjected to two rounds of Heterogeneous refinement (K=6, 5) to remove junk particles. The best class containing 429,738 particles was re-extracted and refined using Non-Uniform (NU) refinement. These particles were subjected to reference-based motion correction and classified in 2D to generate a subset of 428,907 cleaned and polished particles. These particles were subjected to Non-Uniform (NU) refinement and Local refinement, yielding a 3D cryo-EM map at an estimated overall resolution of 3.27 Å. The local resolution was estimated via cryoSPARC.

### Model building and refinement

The LptD/E/M structural model of *E. coli* predicted by AlphaFold2 (Jumper *et al*, 2021; Varadi *et al*, 2024)was docked into the obtained cryo-EM map UCSF Chimera X. The fitted structures were manually adjusted in Coot (Emsley *et al*, 2010). Further refinements were carried out using the phenix.real_space_refine in PHENIX (Adams *et al*, 2010). The data processing and refinement statistics are summarized in Figure S10. The molecular model and cryo-EM map were visualized with PyMOL (https://pymol.org/) or UCSF ChimeraX (Pettersen *et al*, 2021).

### *In vivo* photo-crosslinking analysis

pEVOL-pBpF expresses an evolved tRNA/aminoacyl-tRNA synthetase pair that enables *in vivo* incorporation of *p*BPA into an *amber* codon site of a target protein via *amber* suppression (Young *et al*, 2010). UV exposure of a cell expressing a *p*BPA-incorporated protein induces the formation of covalent crosslinking products between *p*BPA in the target protein and a nearby protein, allowing the detection of their *in vivo* interaction (Chin & Schultz, 2002; Young *et al*, 2010; Miyazaki *et al*, 2020a). Cells of RM4749 (Δ*lptM*) carrying pEVOL-pBpF and pTWV228-*lptM(amb)-his_10_* plasmids were grown at 30 °C in LB medium supplemented with 0.5 mM *p*BPA and 0.02% arabinose until the early log phase and induced with 1 mM IPTG for 1 h. A half volume of the cell cultures was placed on a Petri dish and UV-irradiated at 4 °C for 10 min using a B-100AP UV lamp (365 nm; UVP, LLC.) at a distance of 4 cm. The other half was kept on ice as non-UV-irradiated samples. Total cellular proteins were precipitated with 5% trichloroacetic acid (TCA), washed with acetone, solubilized in SDS sample buffer containing ME and boiled at 98 °C for 5 min. The samples were subjected to SDS‒PAGE and immunoblotting analysis.

### LptMxLptD disulfide crosslinking

Cells of RM5205 (P*_ara_-lptD*, Δ*lptM*) carrying a combination of plasmids encoding WT or a Cys-introduced mutant of LptM-3xFLAG and LptD-His_10_ were inoculated into LB medium containing 0.2% arabinose at 30 °C for 15–16 h. The overnight culture was washed and resuspended in LB medium. The cell suspensions were inoculated into LB medium supplemented with 1 mM IPTG at 30 °C for 3 h. Total cellular proteins were precipitated with 5% TCA, washed with acetone, and suspended in SDS sample buffer containing 12.5 mM NEM to block the free SH groups of the Cys residues. Half of the samples were mixed with 10% 2-mercaptoethanol, boiled at 98 °C for 5 min and analyzed by 7.5% Laemmli SDS‒PAGE and immunoblotting with anti-LptD and anti-FLAG antibodies.

## Data availability

### Author contributions

R.M. conceptualized the study. R.M. and M.K. performed the functional analyses. R.M., M.K., and H.K. performed the cryo-EM analysis. H.K. processed the cryo-EM data. H.K. and T.T. refined the structure model. All of the authors discussed the data. R.M. and T.T. wrote the manuscript and supervised the study.

### Disclosure and competing interests statement

The authors declare that they have no competing interests.

## Acknowledgments

We thank Kayo Abe for secretarial assistance and Kunihito Yoshikaie for technical support. Cryo-EM data collection was performed using a Glacios cryo-TEM with the support of the Cryo-EM Facility, Institute for Life and Medical Sciences, Kyoto University. This work was supported by JSPS/MEXT KAKENHI (Grant Nos. JP21KK0126, JP22K15061, JP22H05567 to R.M., Grant Nos. JP23K14146 to H.K., and Grant Nos. JP23K23850, JP22H02567, JP22H02586, JP21H05155, JP21H05153, JP21K19226, JP21KK0125 to T.T.) and private research foundations (the Institute for Fermentation (Y-2024-02-006) to R.M., and the Chemo-Sero-Therapeutic Research Institute, Takeda Science Foundation to T.T.)

## Notes

### Competing Interest Statement

The authors have declared no competing interest.

