## Supplemental Information for "Structural basis of lipopolysaccharide translocon assembly mediated by the small lipoprotein LptM"

### Figure S1

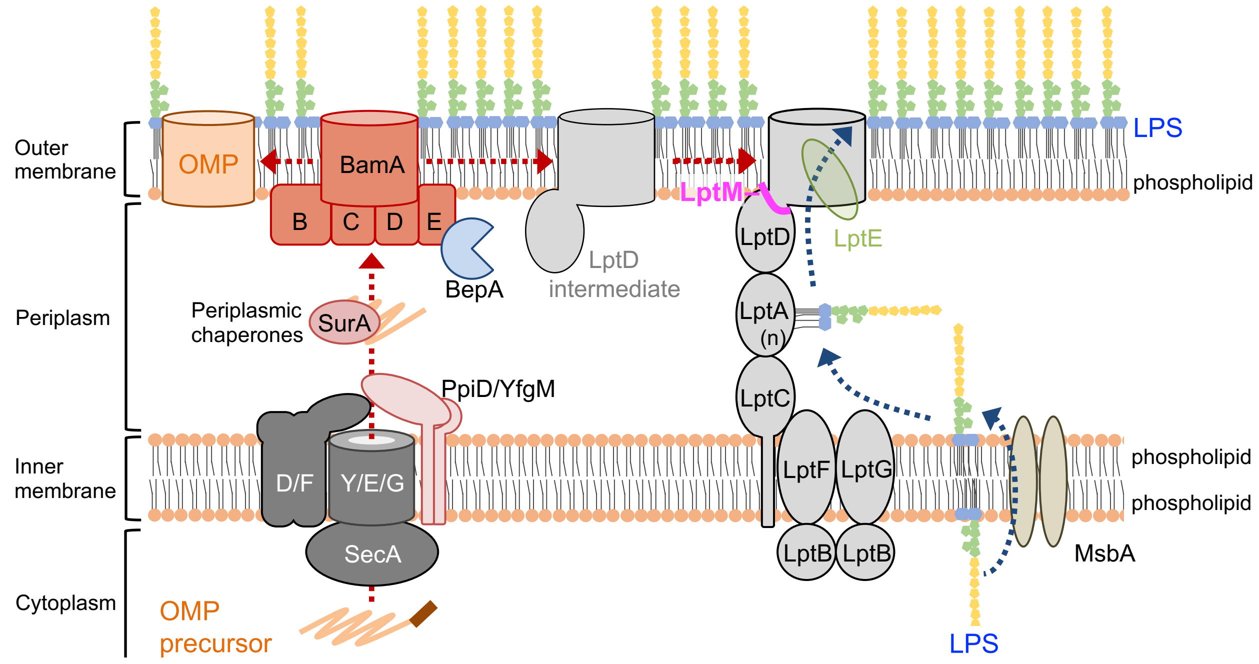


**Figure S1. Schematic representation of the BAM complex, the Lpt complex and related factors on the cell surface of *E. coli*.**

### Figure S2

A

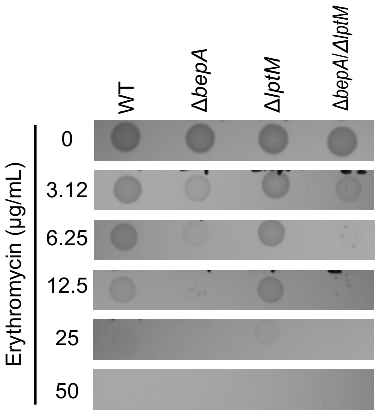


B

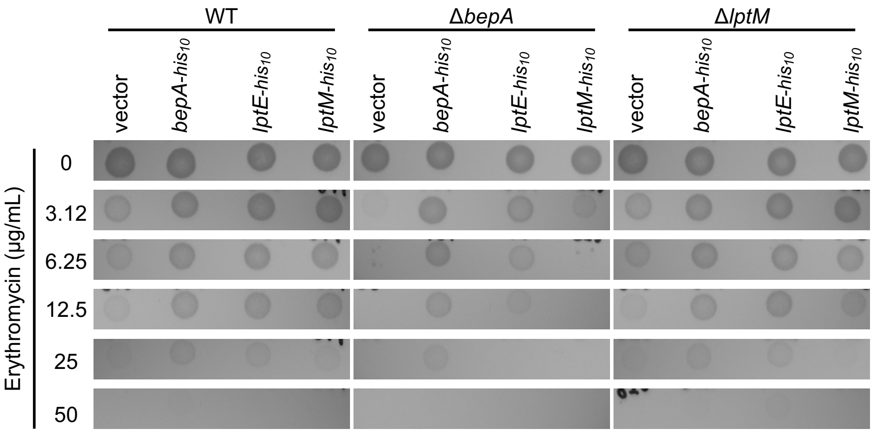


**Figure S2. The original data of the erythromycin sensitivity assay**
(**A**) The original data of Figure 1C. (**B**) The original data of Figure 1E.

### Figure S3

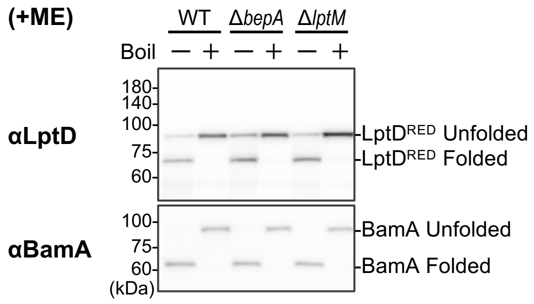


**Figure S3. Heat modifiability of the LptD intermediate in Δ*bepA* and Δ*lptM* mutant cells on SDS–PAGE.**
The same samples shown in Figure 1F were analyzed by 7.5% Laemmli SDS–PAGE under reducing (+ME) conditions at 4 ºC and immunoblotting with the indicated antibodies.

### Figure S4

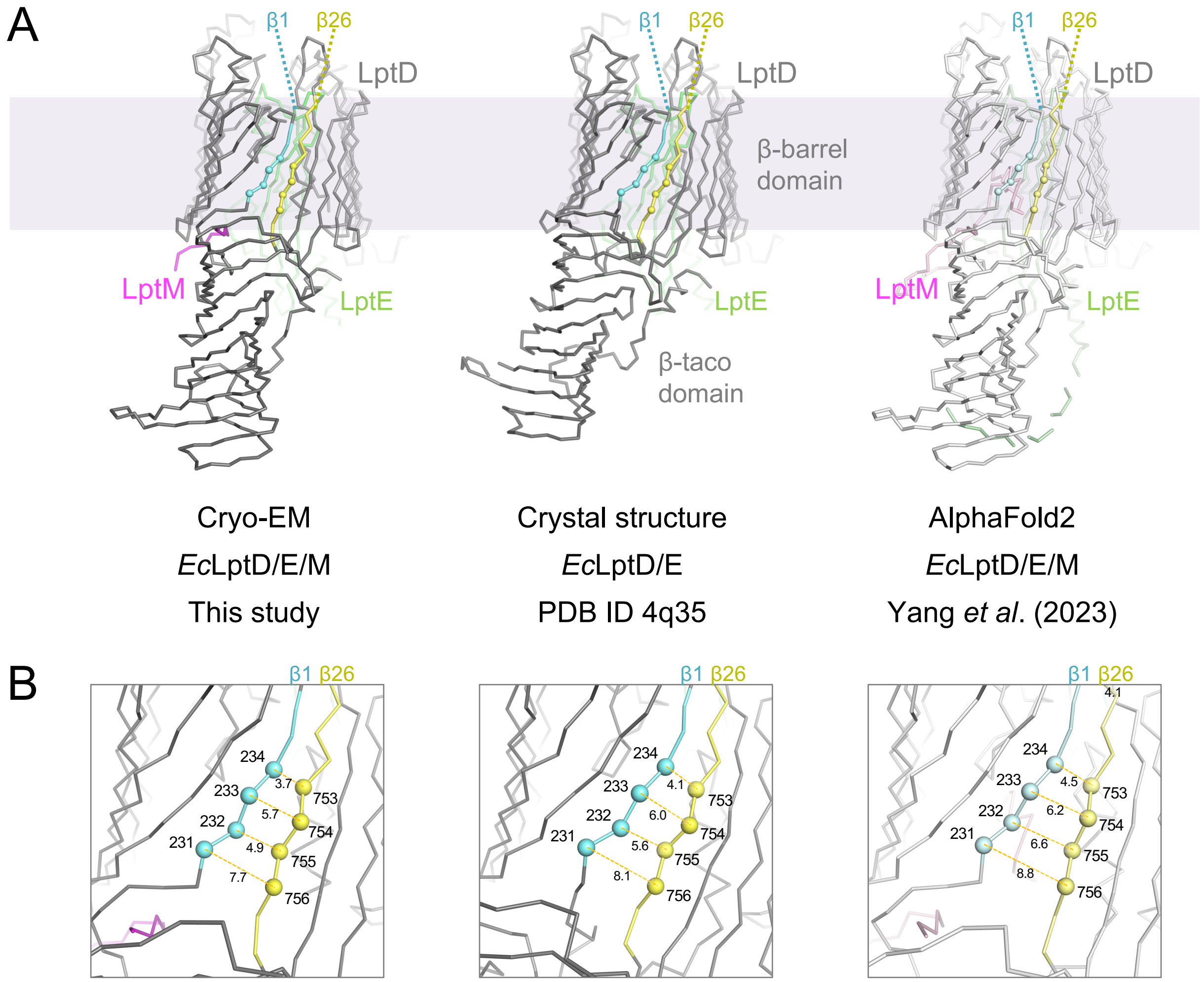


**Figure S4. Structures of the LptD/E complex.**
(**A**) Overall structures of the LptD/E complex. Left: The cryo-EM structure of *E. coli* LptD/E/M. Center: Crystal structure of *E. coli* LptD/E (PDB ID: 4q35). Right: AlphaFold2 model of *E. coli* LptD/E/M reported by Yang *et al.* (2023). LptD, LptM, and LptE are shown in gray, magenta, and green, respectively. β1 and β26 of LptD are highlighted in cyan and yellow, respectively. (**B**) Close-up views of Panel A. Distances between the indicated Cα atoms are displayed.

### Figure S5

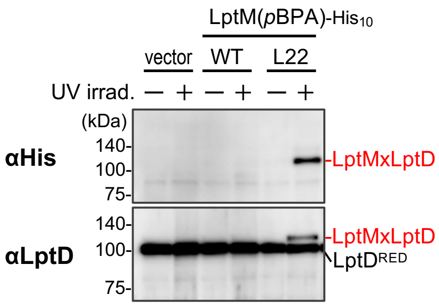


**Figure S5. *In vivo* photo-crosslinking analysis of LptM L22*p*BPA**
RM4749 (Δ*lptM*) cells harboring pEVOL-pBpF and pTWV228 or pTWV228-*lptM(amb)-his*$_{10}$ plasmids were grown and analyzed via SDS–PAGE and immunoblotting, as shown in Figure 4D.

### Figure S6

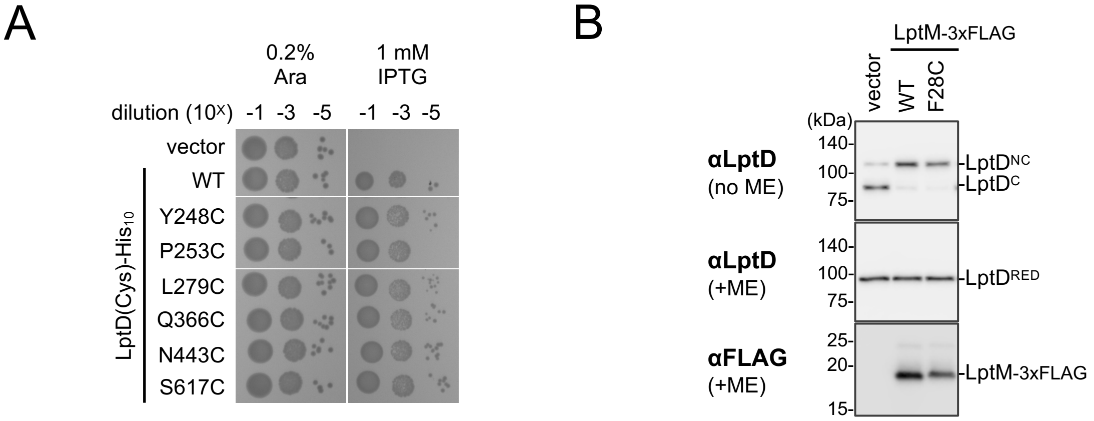


**Figure S6. Activities of LptD Cys mutants and the LptM F28C mutant used for LptMxLptD disulfide crosslinking**
(**A**) Complementation activity of the LptD mutants with an engineered Cys residue. Cells of RM3588 ($P_{ara}$-*lptD*) carrying pTWV228 or pTWV228-*lptD(Cys)-his$_{10}$* plasmids were grown at 30 ºC in LB medium supplemented with 0.2% arabinose for 2.5 h. The cells were subsequently washed, suspended in saline, and serially diluted with saline (to approximately $10^9$ cells/mL). A total of 2.5 µL of each of the diluted cells was spotted on an LB agar plate containing 0.2% arabinose (positive control) or 1 mM IPTG (for the induction of LptD(Cys) from a plasmid). The plates were incubated at 30 ºC for 22 h. (**B**) Activity of the LptM F28C mutant. Cells of RM4749 (Δ*lptM*) carrying pSTD689 or pSTD689-*lptM(Cys)-3xflag* plasmids were grown and analyzed via SDS–PAGE and immunoblotting, as shown in Figure 2B.

### Figure S7

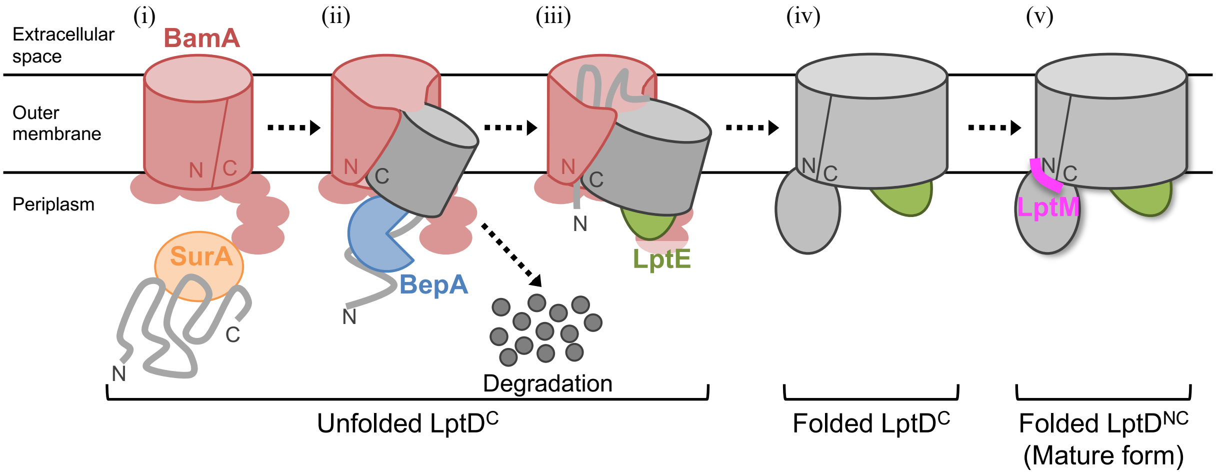


**Figure S7. Model for the proposed LptD assembly process**

### Figure S8

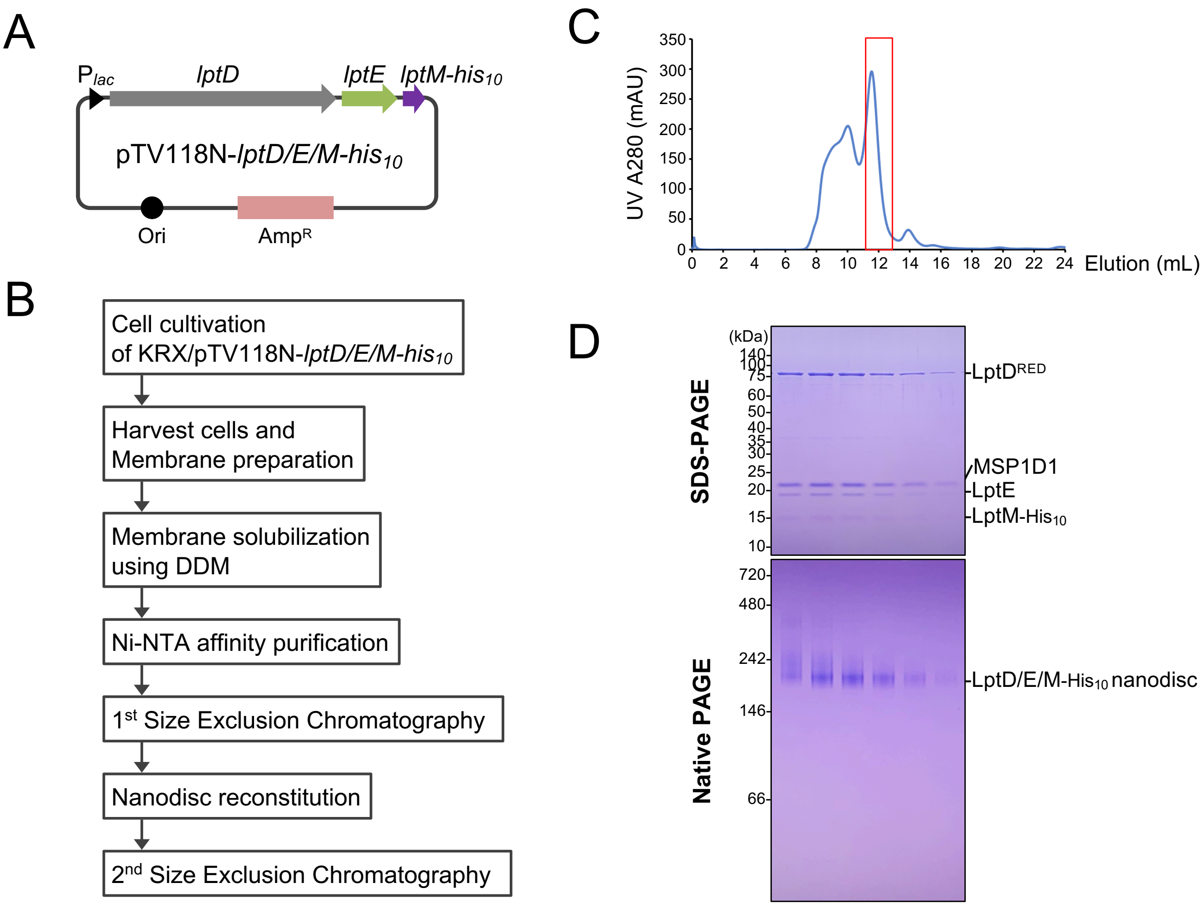


**Figure S8. Preparation of the nanodisc reconstituted EcLptD/E/M complex**
(**A**) Schematic of a plasmid encoding *lptD/lptE/lptM-his*$_{10}$. (**B**) Workflow for purification and nanodisc reconstitution of the *Ec*LptD/E/M complex. (**C**) Size-exclusion chromatogram of the nanodisc-reconstituted *Ec*LptD/E/M complex. (**D**) SDS–PAGE and Native–PAGE gels showing peak fractions from size-exclusion chromatography of the complex in **C**.

### Figure S9

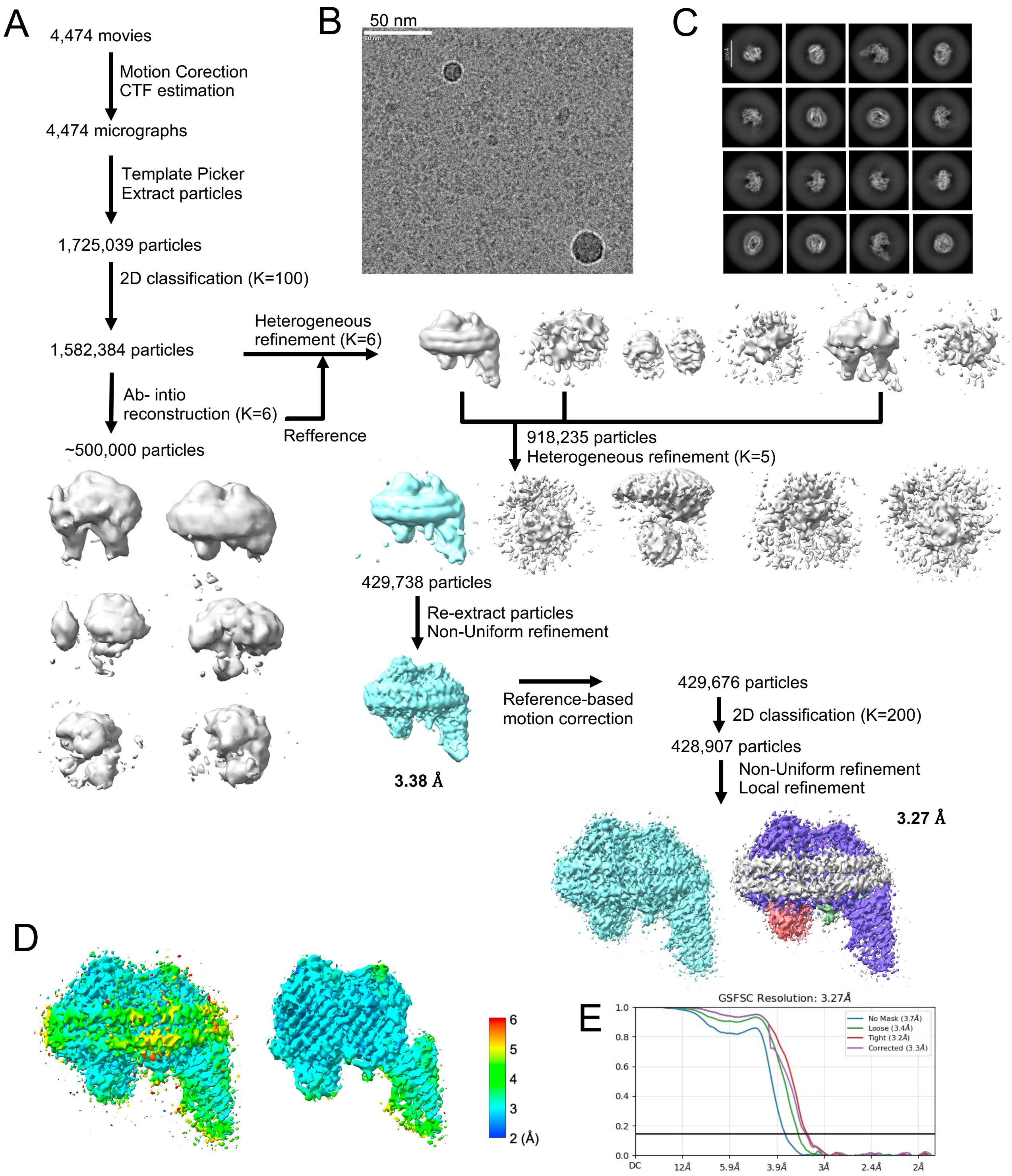


**Figure S9. Cryo-EM data-processing workflow for the *Ec*LptD/E/M complex**
(**A**) Schematic of the preprocessing, classification, and refinement procedures. (**B**) Representative raw electron micrograph of the nanodisc-reconstituted *Ec*LptD/E/M complex. (**C**) Representative 2D classes. (**D**) Final reconstruction colored according to the local resolution. (**E**) Gold-standard Fourier shell correlation (FSC) curve.

### Figure S10

| | *Ec*LptDEM-ND<br>EMD-62452, PDB ID: 9kn3 |
|---|---|
| Data collection and processing | |
| Magnification | x150,000 |
| Voltage (kV) | 200 |
| Electron exposure (e−/Å$^2$) | 50 |
| Defocus range (μm) | -0.6 to -1.6 |
| Pixel size (Å) | 0.924 |
| Symmetry imposed | C1 |
| Initial particle images (no.) | 1,725,039 |
| Final particle images (no.) | 428,907 |
| Map resolution (Å) | 3.27 |
| FSC threshold | 0.143 |
| Map resolution range (Å) | 9.99 to 2.02 |
| Refinement | |
| Initial model used (PDB code) | AlphaFold model |
| Model composition | |
| Non-hydrogen atoms | 7392 |
| Protein residues | 917 |
| Ligands | 0 |
| B factors (Å$^2$) | |
| Protein | 143.5 |
| Ligand | 0 |
| R.m.s. deviations | |
| Bond lengths (Å) | 0.009 |
| Bond angles (°) | 0.824 |
| Validation | |
| MolProbity score | 2.03 |
| Clashscore | 8.05 |
| Poor rotamer (%) | 0.5 |
| Ramachandran plot | |
| Favored (%) | 88.23 |
| Allowed (%) | 11.66 |
| Disallowed (%) | 0.11 |

**Figure S10. Cryo-EM data collection, refinement and statistics.**

**Table S1. Strains used in this study.**

| Strain | Genotype | Reference |
|---|---|---|
| AD16 | Δ*pro-lac thi*/F' $lacI^{q}$ *Z*Δ*M15* $Y^{+}$ $pro^{+}$ | Kihara *et al*., 1995 |
| SN56 | AD16, Δ*bepA* | Narita *et al*., 2013 |
| RM4749 | AD16, Δ*lptM*::*kan* | This study |
| RM4752 | AD16, Δ*bepA* Δ*lptM*::*kan* | This study |
| MC4100 | $F^{-}$ *araD139* Δ*(argF-lac)U169 rpsL150 relA1 flbB5301 deoC1 ptsF25 rbsR* | Silhavy *et al*., 1984 |
| CU141 | MC4100/F' $lacI^{q}$ $lacZ^{+}$, $Y^{+}$, $A^{+}$ | Akiyama *et al*., 1994 |
| HM1742 | CU141, $ara^{+}$ | Mori and Ito, 2006 |
| RM3588 | HM1742, *kan araC*-$P_{araBAD}$-*lptD* | Miyazaki *et al*., 2021 |
| RM5181 | HM1742, *araC*-$P_{araBAD}$-*lptD* | This study |
| RM5205 | HM1742, *araC*-$P_{araBAD}$-*lptD* Δ*lptM*::*kan* | This study |
| DY330 | W3110, Δ*lacU169 gal490* λ*cI857* Δ(*cro-bioA*) | Yu e*t al*., 2000 |
| RM4717 | DY330, Δ*lptM*::*kan* | This study |
| KRX | Δ*ompT endA1 recA1 gyrA96*($Nal^{R}$) *thi-1 hsdR17*($r_K^{-}$, $m_K^{+}$), $e14^{-}$ ($McrA^{-}$) *relA1 supE44* Δ(*lac-pro*AB) Δ(*rha*BAD)::T7 RNA polymerase / F′[*traD36* Δ*ompP* $proA^{+}B^{+}$ $lacI^{q}$ Δ*lac*ZM15] | Promega |

**Table S2. Plasmids used in this study.**

| Plasmid | Vector | Encoded gene and description | Reference or source |
|---|---|---|---|
| pEVOL-pBpF | | p15A-derivative encoding an evolved *M. jannaschii* aminoacyl-tRNA synthetase/suppressor tRNA pair for incorporation of *p*BPA; $Cm^R$ | Young *et al*., 2010 |
| pTWV228 | | Expression vector; $P_{lac}$, $Amp^R$ | Takara Bio |
| pRM1441 | pTWV228 | *lptM-his$_{10}$* | This study |
| pRM1442 | pTWV228 | *lptM(L22amb)-his$_{10}$* | This study |
| pRM1443 | pTWV228 | *lptM(K23amb)-his$_{10}$* | This study |
| pRM1444 | pTWV228 | *lptM(P25amb)-his$_{10}$* | This study |
| pRM1445 | pTWV228 | *lptM(Y27amb)-his$_{10}$* | This study |
| pRM1446 | pTWV228 | *lptM(P29amb)-his$_{10}$* | This study |
| pRM1447 | pTWV228 | *lptM(A31amb)-his$_{10}$* | This study |
| pRM1448 | pTWV228 | *lptM(K33amb)-his$_{10}$* | This study |
| pRM1449 | pTWV228 | *lptM(A35amb)-his$_{10}$* | This study |
| pRM1450 | pTWV228 | *lptM(P37amb)-his$_{10}$* | This study |
| pRM1451 | pTWV228 | *lptM(T39amb)-his$_{10}$* | This study |
| pRM1452 | pTWV228 | *lptM(P41amb)-his$_{10}$* | This study |
| pRM1453 | pTWV228 | *lptM(E43amb)-his$_{10}$* | This study |
| pRM1454 | pTWV228 | *lptM(Q45amb)-his$_{10}$* | This study |
| pRM1455 | pTWV228 | *lptM(Q47amb)-his$_{10}$* | This study |
| pRM1456 | pTWV228 | *lptM(T49amb)-his$_{10}$* | This study |
| pRM1457 | pTWV228 | *lptM(P51amb)-his$_{10}$* | This study |
| pRM1458 | pTWV228 | *lptM(K53amb)-his$_{10}$* | This study |
| pRM1459 | pTWV228 | *lptM(N54amb)-his$_{10}$* | This study |
| pRM1460 | pTWV228 | *lptM(D55amb)-his$_{10}$* | This study |
| pRM1461 | pTWV228 | *lptM(R56amb)-his$_{10}$* | This study |
| pRM1462 | pTWV228 | *lptM(A57amb)-his$_{10}$* | This study |
| pRM1463 | pTWV228 | *lptM(G59amb)-his$_{10}$* | This study |
| pRM1464 | pTWV228 | *lptM(G61amb)-his$_{10}$* | This study |
| pRM1465 | pTWV228 | *lptM(P62amb)-his$_{10}$* | This study |
| pRM1466 | pTWV228 | *lptM(S63amb)-his$_{10}$* | This study |
| pRM1467 | pTWV228 | *lptM(Q64amb)-his$_{10}$* | This study |
| pRM1468 | pTWV228 | *lptM(V65amb)-his$_{10}$* | This study |
| pRM1469 | pTWV228 | *lptM(Y67amb)-his$_{10}$* | This study |
| pRM1470 | pTWV228 | *lptM* | This study |

| | | | |
|---|---|---|---|
| pRM1531 | pTWV228 | *lptM(G21A)-his$_{10}$* | This study |
| pRM1532 | pTWV228 | *lptM(L22A)-his$_{10}$* | This study |
| pRM1533 | pTWV228 | *lptM(K23A)-his$_{10}$* | This study |
| pRM1534 | pTWV228 | *lptM(G24A)-his$_{10}$* | This study |
| pRM1535 | pTWV228 | *lptM(P25A)-his$_{10}$* | This study |
| pRM1536 | pTWV228 | *lptM(L26A)-his$_{10}$* | This study |
| pRM1537 | pTWV228 | *lptM(Y27A)-his$_{10}$* | This study |
| pRM1538 | pTWV228 | *lptM(F28A)-his$_{10}$* | This study |
| pRM1539 | pTWV228 | *lptM(G21C)-his$_{10}$* | This study |
| pRM1540 | pTWV228 | *lptM(L22C)-his$_{10}$* | This study |
| pRM1541 | pTWV228 | *lptM(K23C)-his$_{10}$* | This study |
| pRM1542 | pTWV228 | *lptM(G24C)-his$_{10}$* | This study |
| pRM1543 | pTWV228 | *lptM(P25C)-his$_{10}$* | This study |
| pRM1544 | pTWV228 | *lptM(L26C)-his$_{10}$* | This study |
| pRM1545 | pTWV228 | *lptM(Y27C)-his$_{10}$* | This study |
| pRM1546 | pTWV228 | *lptM(F28C)-his$_{10}$* | This study |
| pRM1547 | pTWV228 | *lptM(G21W)-his$_{10}$* | This study |
| pRM1548 | pTWV228 | *lptM(L22W)-his$_{10}$* | This study |
| pRM1549 | pTWV228 | *lptM(K23W)-his$_{10}$* | This study |
| pRM1550 | pTWV228 | *lptM(G24W)-his$_{10}$* | This study |
| pRM1551 | pTWV228 | *lptM(P25W)-his$_{10}$* | This study |
| pRM1552 | pTWV228 | *lptM(L26W)-his$_{10}$* | This study |
| pRM1553 | pTWV228 | *lptM(Y27W)-his$_{10}$* | This study |
| pRM1554 | pTWV228 | *lptM(F28W)-his$_{10}$* | This study |
| pRM1601 | pTWV228 | *lptM(G21W, L22amb)-his$_{10}$* | This study |
| pRM1604 | pTWV228 | *lptM(G21W, K53amb)-his$_{10}$* | This study |
| pRM267 | pTWV228 | *lptD-his$_{10}$* | Daimon *et al.*, 2017 |
| pRM1577 | pTWV228 | *lptD(Y248C)-his$_{10}$* | This study |
| pRM1578 | pTWV228 | *lptD(P253C)-his$_{10}$* | This study |
| pRM1580 | pTWV228 | *lptD(L279C)-his$_{10}$* | This study |
| pRM1581 | pTWV228 | *lptD(Q366C)-his$_{10}$* | This study |
| pRM1582 | pTWV228 | *lptD(N443C)-his$_{10}$* | This study |
| pRM1583 | pTWV228 | *lptD(S617C)-his$_{10}$* | This study |
| pSTD689 | | Expression vector; $P_{lac}$, $Spc^R$ | Kanehara *et al.*, 2003 |
| pRM867 | pSTD689 | *bepA-his$_{10}$* | This study |
| pRM1381 | pSTD689 | *lptE-his$_{10}$* | This study |

| | | | |
|---|---|---|---|
| pRM1383 | pSTD689 | *lptM-his$_{10}$* | This study |
| pRM1555 | pSTD689 | *lptM-3xflag* | This study |
| pRM1566 | pSTD689 | *lptM(F28C)-3xflag* | This study |
| pTV118N | | Expression vector; $P_{lac}$, $Amp^R$ | Takara Bio |
| pRM1557 | pTV118N | *lptD/lptM-his$_{10}$* | This study |
| pRM1606 | pTV118N | *lptD/lptE/lptM-his$_{10}$* | This study |

**Table S3. Primers used in this study.**

| Name | Sequence (5’ to 3’) |
|---|---|
| d-lptM-4F | ATAGAAAGCAGAAAGCGATGAACTTTACAGGCAATCCATAGTGTAGGCTGGAGCTGCTTC |
| d-lptM-4R | GCATCATTTACTCCAATCACGCGGGTACAGAAACTGACTTCATATGAATATCCTCCTTA |
| pSTD-H-F | CTCGAGCATCATCATCATC |
| pSTD-H-R | CATAACCTCTATCCTGTATTTC |
| lptE-F | AGGATAGAGGTTATGCGATATCTGGCAACATTG |
| lptE-R | ATGATGATGCTCGAGGTTACCCAGCGTGGTGGAG |
| lptM-F | AGGATAGAGGTTATGAAAAACGTGTTTAAGGC |
| lptM-R | ATGATGATGCTCGAGGTAATTCACCTGGGATGGAC |
| pTV-F | GGCACTGGCCGTCGTTTTAC |
| pTV-R | CATGGTCTGTTTCCTGTGTG |
| lptD-F | AGGAAACAGACCATGATGAAAAAACGTATCCCCAC |
| lptD-R | ACCCCTCCGCCGGCCGCTGCTCACAAAGTGTTTTGATACG |
| lptE-F2 | GGCCGGCGGAGGGGTGCGATATCTGGCAAC |
| lptE-R2 | TCCGCCGGCCGCTGCTTAGTTACCCAGCGTGGTGG |
| lptM-F2 | GCAGCGGCCGGCGGAGGGATGAAAAACGTGTTTAAGGCAC |
| lptM-R2 | ACGACGGCCAGTGCCTTAATGATGATGATGATGATG |
| plptDM-F | ATGAAAAACGTGTTTAAGGCAC |
| plptDM-R | TCACAAAGTGTTTTGATACGG |
| LptE-F3 | ACACTTTGTTTGTGAGCAGCGGCCGGCGGAGGGGTGCGATATCTGGCAACATTG |
| LptE-R3 | AAACACGTTTTTCATCCCTCCGCCGGCCGCTGCTTAGTTACCCAGCGTGGTGG |
